# JEDI: Circular RNA Prediction based on Junction Encoders and Deep Interaction among Splice Sites

**DOI:** 10.1101/2020.02.03.932038

**Authors:** Jyun-Yu Jiang, Chelsea J.-T. Ju, Junheng Hao, Muhao Chen, Wei Wang

## Abstract

Circular RNA is a novel class of endogenous non-coding RNAs that have been largely discovered in eukaryotic transcriptome. The circular structure arises from a non-canonical splicing process, where the donor site backsplices to an upstream acceptor site. These circular form of RNAs are conserved across species, and often show tissue or cell-specific expression. Emerging evidences have suggested its vital roles in gene regulation, which are further associated with various types of diseases. As the fundamental effort to elucidate its function and mechanism, numerous efforts have been devoted to predicting circular RNA from its primary sequence. However, statistical learning methods are constrained by the information presented with explicit features, and the existing deep learning approach falls short on fully exploring the positional information of the splice sites and their deep interaction.

We present an effective and robust end-to-end framework, JEDI, for circular RNA prediction using only the nucleotide sequence. Our framework first leverages the attention mechanism to encode each junction site based on deep bidirectional recurrent neural networks and then presents the novel cross-attention layer to model deep interaction among these sites for backsplicing. Finally, JEDI is capable of not only addressing the task of circular RNA prediction but also interpreting the relationships among splice sites to discover the hotspots for backsplicing within a gene region. Experimental evaluations demonstrate that JEDI significantly outperforms several state-of-the-art approaches in circular RNA prediction on both isoform-level and gene-level. Moreover, JEDI also shows promising results on zero-shot backsplicing discovery, where none of the existing approaches can achieve.

The implementation of our framework is available at https://github.com/hallogameboy/JEDI.

## 1 Introduction

As a special type of long non-coding RNA (lncRNA), circular RNA (circRNA) has received rising attention due to its circularity and implications in a myriad of diseases, such as cancer and Alzheimer’s [9, 39]. It arises during the process of alternative splicing of protein-coding genes, where the 5′ end of an exon is covalently ligated to the 3′ end of the same exon or a downstream exon, forming a closed continuous loop structure. This mechanism is also known as “backsplicing.” The circular structure provides several beneficial properties over the linear RNAs. To be more specific, it can serve as templates for rolling circle amplification of RNAs [5], rearrange the order of genetic information [30], resistant to exonuclease-mediated degradation [27], and create a constraint on RNA folding [30]. Although the consensus of biological functions, mechanisms, and biogenesis remains unclear for most circRNAs [4, 50], there are emerging studies suggesting their roles in acting as sponges for microRNAs [20, 34], RNA-binding protein competition [2], and inducing host gene transcription [33]. Evidently, as a fundamental step to facilitate the exploration of circRNA, it is essential to have a high-throughput approach to identify the circRNAs.

Multiple factors can contribute to the formation of circRNAs. These factors include complementary sequences in flanking introns [25], the presence of inverted repeats [10], number of ALU and tandem repeats [27], and SNP density [45]. These factors, together with the evolution conservation and secondary structure of RNA molecules, have been considered as the discriminative features for circRNA identification. Several research efforts [7, 37, 47] have leveraged these features to train a conventional statistical learning model to distinguish circRNAs from other lncRNAs. These statistical learning algorithms include support vector machines (SVM), random forest (RF), and multi-kernel learning. However, methods along this line often require an extensive domain-specific feature engineering process. Moreover, the selected features may not provide sufficient coverage to characterize the backsplicing event.

Recently, the rising of deep learning architectures have been widely adopted as an alternative learning algorithm that can alleviate the inadequacy of conventional statistical learning methods. Specifically, these deep learning algorithms provide powerful functionality to process large-scale data and automatically extract useful features for object tasks [31]. In the domain of circRNA prediction, the convolution neural network (CNN) is the architecture that has been widely explored to automatically learn the important features for prediction, either from the primary sequence [6, 48] or secondary structure [11]. Although CNN is capable of capturing important local patterns on gene sequences for prediction, positional information and global context of each splice site cannot be recognized. One of these approaches [6] attempts to address this issue by applying recurrent neural networks (RNNs) to learn sequential and contextual information; however, the essential knowledge, such as splice sites and junctions, are still ignored.

Understanding the properties of splice sites and their relationships can be one of the keys to master RNA splicing and the formation of circular RNAs because the splicing event can be considered as interaction among those splice sites. To fathom the relations between splice sites, circDeep [6] explicitly matches the splices sites on the nucleotide level for predicting circular RNAs. DeepCirCode [48] utilizes CNNs to model the flanking regions around two splice sites to identify if there is a backsplice. However, all of the existing methods fail in modeling deep interaction among splice sites for circular RNA prediction. For example, circDeep only measures shallow interaction among splice sites on the nucleotide level; DeepCirCode can only tackle a single pair of splice sites for backsplicing prediction without the capacity of modeling more complex relations among splice sites on multi-isoform genes. Hence, there is still a huge room for improvement on the way to comprehensively understand splice sites and their interaction about backsplicing and the formation of circular RNAs.

In this paper, the framework of Junction Encoder with Deep Interaction (JEDI) is proposed to address the limitations in circular RNA prediction. More precisely, we focus on predicting the existence of circular RNAs from either the reference gene/isoform sequences or assembled transcript sequences by modeling splice sites and their deep interaction with deep learning techniques. First, the attentive junction encoders are presented to derive continuous embedding vectors for acceptor and donor splice sites based on their flanking regions around junctions. Based on the acceptor and donor embeddings, we propose the novel cross-attention layer to model deep interaction between acceptor and donor sites, thereby inferring cross-attentive embedding vectors. Finally, the attention mechanism is applied to determine acceptors and donors that are more important than other ones to predict if there is a circRNA. It is also important to note that the interpretability of the attention mechanism and the cross-attention layer enables JEDI to automatically discover backsplicing without training on any annotated backspliced sites.

Our contributions are three-fold. First, to the best of our knowledge, this work is the first study to model deep interaction among splice sites for circular RNA prediction. The more profound understandings of the relationships among splice sites can intuitively benefit circular RNA prediction in implying backsplicing. Second, we propose a robust and effective end-to-end framework JEDI to deal with both isoform-level and gene-level circular RNA prediction based on the attention mechanism and the innovative cross-attention layer. More specifically, JEDI is capable of not only deriving appropriate representations from junction encoders but also routing the importance about forming circular RNAs on different levels. Third, JEDI creates a new opportunity of transferring the knowledge from circular RNA prediction to backsplicing discovery based on its extensive usage of attention mechanisms. Extensive experiments on human circRNAs have demonstrated that JEDI significantly outperforms eight competitive baseline methods on both isoform-level and gene-level. The independent study on mouse circRNAs also indicates that JEDI is robust to transfer knowledge for circular RNA prediction from human data to mouse data. In addition, we conduct the experiments to demonstrate that JEDI can automatically discover backspliced site pairs without any further annotations. Finally, an in-depth analysis on model hyper-parameters and run-time presents the robustness and efficiency of JEDI.

## 2 Related Work

Current works to discover circular RNA can be divided into two categories: one relies on detecting back-spliced junction reads from RNA-Seq data; the other examines features directly from transcript sequences.

The first category aims at detecting circRNA from expression data. It is mainly achieved by searching for chimeric reads that join the 3′-end to the upstream 5′-end with respect to a transcript sequence [4]. Existing algorithms include **MapSplice** [49], **CIRCexplorer** [51], **KNIFE** [44], **findcirc** [34], and **CIRI** [13, 14]. These algorithms can be quite sensitive to the expression abundance, as circRNAs are often lowly expressed and fail to be captured with low sequencing coverage [4]. In the comparison conducted by Hansen et al. [21], the findings suggest dramatic differences among these algorithms in terms of sensitivity and specificity. Other caveats are reflected in long duration, high RAM usage, and/or complicated pipeline.

The second category focuses on predicting the circRNA based on transcript sequences. Methods in this category leverage different features and learning algorithms to distinguish circRNA from other lncRNAs. **PredicircRNA** [37] and **H-ELM** [7] develop different strategies to extract discriminative features, and employ conventional statistical learning algorithms, i.e. multiple kernel learning for PredicircRNA and hierarchical extreme learning machine for H-ELM, to build a classifier. Statistical learning approaches require explicit feature engineering and selection. However, the extracted features are dedicated to specific facets of the sequence properties and present a limited coverage on the interaction information between the donor and acceptor sites. **circDeep** [6] and **DeepCirCode** [48] are two pioneering methods that employ deep learning architectures to automatically learn complex patterns from the raw sequence without extensive feature engineering. circDeep uses convolution neural networks (CNNs) with the bi-directional long short term memory network (Bi-LSTM) to encode the entire sequence, whereas DeepCirCode uses CNNs with maxpooling to capture only the flanking sequences of the back-splicing sites. Although circDeep has claimed to be an end-to-end framework, it requires external resources and strategies to capture the reverse complement matching (RCM) features at the flanking sequence and the conservation level of the sequence. In addition, the RCM features only measure the match scores between sites on the nucleotide-level, and neglect the complicated interaction between two sites. CNNs with max-pooling aim at preserving important local patterns within the flanking sequences. As a result, DeepCirCode fails to retain the positional information of nucleotides and their corresponding convoluted results.

Besides sequence information, a few conventional lncRNA prediction methods also present the potential of discovering circRNA through the secondary structure. **nRC** [11] extracts features from the secondary structures of non-conding RNAs and adopts CNNs framework to classify different types of non-coding RNA. **lncFinder** [19] integrates both the sequence composition and structural information as features and employs random forests. The learning process can be further optimized to predict different types of lncRNA. Nevertheless, none of these methods factor in the information specific to the formation of circRNAs, particularly the interaction information between splicing sites.

## 3 Materials and Methods

In this section, we first formally define the objective of this paper, and then present our proposed deep learning framework, Junction Encoder with Deep Interaction (JEDI), to predict circRNAs.

### 3.1 Preliminary and Problem Statement

Thoecabulary of four nucleotides is denoted as 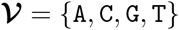. For a gene sequence 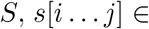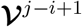 indicates the subsequence from the *i*-th to the *j*-th nucleotide of a sequence *S*. For a gene or an RNA isoform with the sequence *S*, *ε*(*S*) = {(*a*_*i*_, *d*_*i*_)} represents the given exons in the gene or the isoform, where *a*_*i*_ and *d*_*i*_ are the indices of the acceptor and donor junctions of the *i*-th exon in *S*. Using only sequence information, the two goals of this work are listed as follows:

1. **Isoform-level Circular RNA Prediction:** Given a gene sequence *S* and the splicing information of an isoform *ε*(*s*), the goal is to identify whether this RNA isoform is a circRNA.
2. **Gene-level Circular RNA Prediction:** Given a gene sequence *S* and all of its exon-intron boundaries *ε*(*S*), this task aims at predicting if any of the junction pairs can backsplice to form a circRNA.

### 3.2 Framework Overview

Figure 1 illustrates the general schema of JEDI to predict circRNAs. Each acceptor *a*_*i*_ and donor *d*_*j*_ in the gene sequence are first represented by flanking regions *A*_*i*_ and *D*_*i*_ around exon-intron junctions. Two attentive junction encoders then derive embedding vectors of acceptors and donors, respectively. Based on the embedding vectors, we apply the cross-attention mechanism to consider deep interactions between acceptors and donors, thereby obtaining donor-aware acceptor embeddings and acceptor-aware donor embeddings. Finally, the attention mechanism is applied again to learn the provided acceptor and donor representations so that the prediction can be inferred by a fully-connected layer based on the representations.

**Fig. 1:**
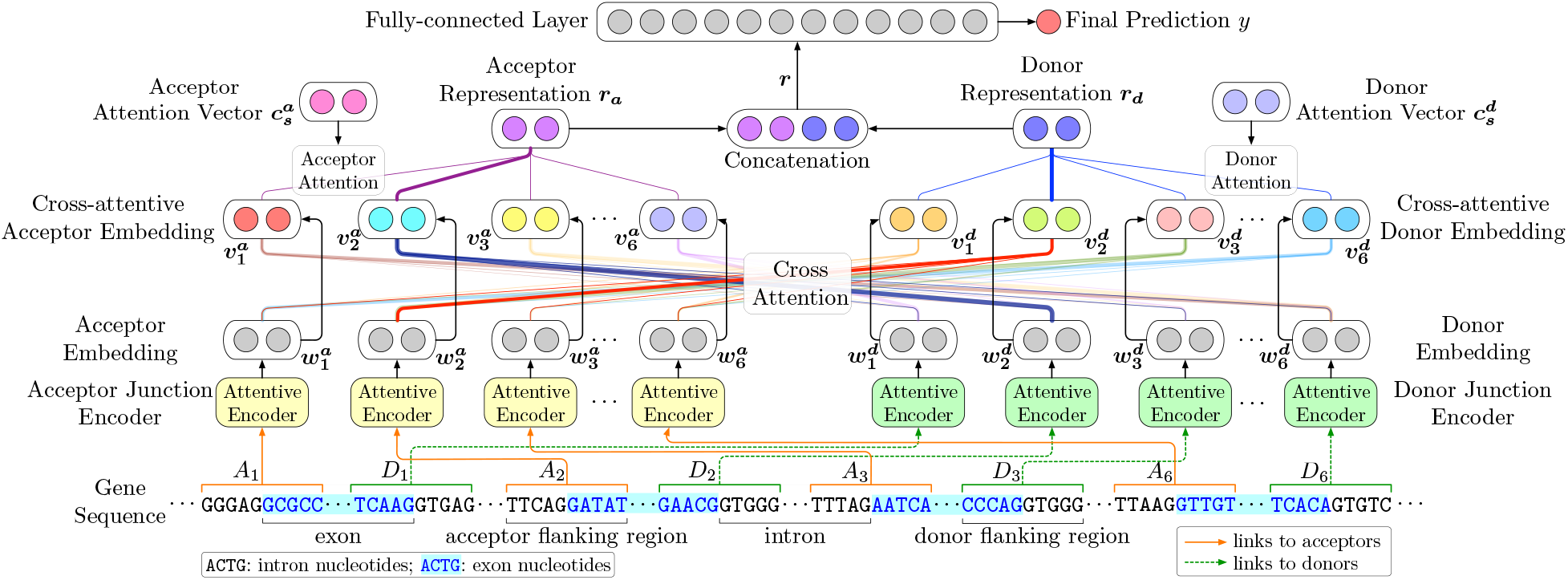
The schema of the proposed framework, *Junction Encoder with Deep Interaction* (JEDI), using the gene `NM_001080433` with six exons as an example, where the second exon forms backsplicing. *A*_*i*_ and *D*_*j*_ represent the *i*-th and *j*-th potential acceptors and donors.

### 3.3 Attentive Junction Encoders

To represent the properties of acceptors and donors in the gene sequence *S*, we utilize the flanking regions around junctions to derive informative embedding vectors. Specifically, as shown in Figure 2, we propose attentive junction encoders using recurrent neural networks (RNNs) and the attention mechanism based on acceptor and donor flanking regions.

**Fig. 2:**
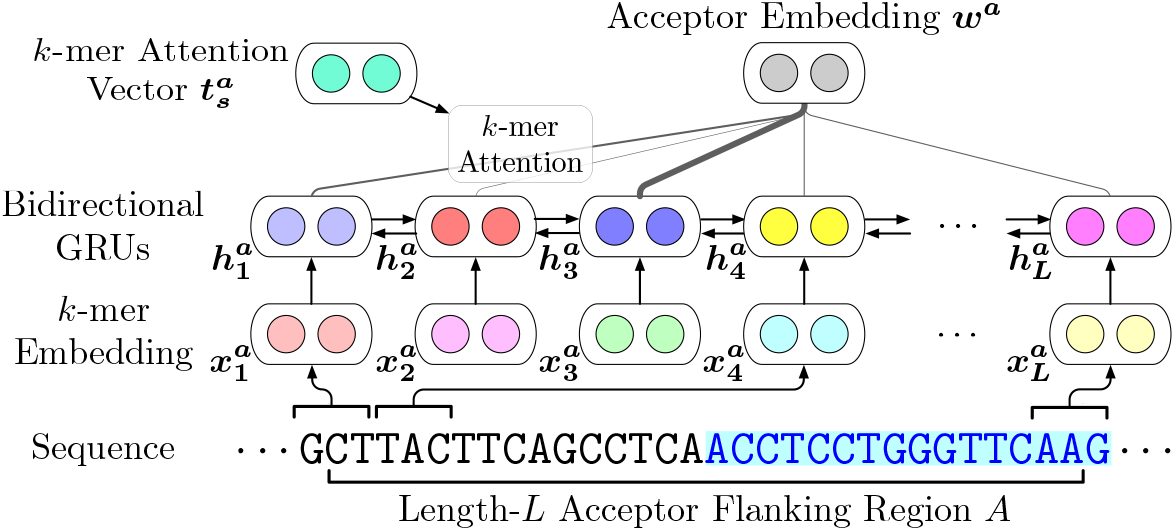
The illustration of the attentive encoder for acceptor junctions. Note that the donor junction encoder shares the same model structure with different model parameters.

#### Flanking Regions as Inputs

For each exon (*a*_*i*_, *d*_*i*_) ∈ *ε*(*S*), length-*L* acceptor and donor flanking regions *A*_*i*_ and *D*_*i*_ can be computed as:

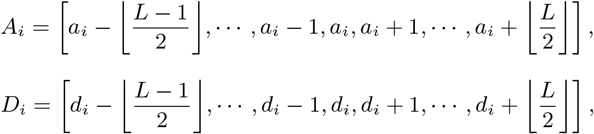

where *A*_*i*_[*j*] and *D*_*i*_[*k*] denote the *j*-th and *k*-th positions on *S* for the flanking regions of the acceptor *a*_*i*_ and the donor *d*_*i*_; the region length *L* is a tunable hyper-parameter.

Suppose we are encoding an acceptor *a* and a donor *d* with the flanking regions *A* and *D* in the gene sequence *S* for the simplicity.

#### *k*-mer Embedding

To represent different positions in the sequence, we use *k*-mers as representations because *k*-mers are capable of preserving more complicated local contexts [29]. Each unique *k*-mer are then mapped to a continuous embedding vector as various deep learning approaches in bioinformatics [6, 35]. Formally, for each position *A*[*j*] and *D*[*k*], the corresponding *k*-mer embedding vectors 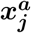 and 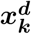 can be derived as follows:

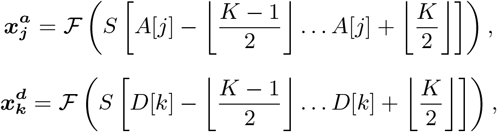

where 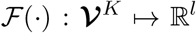 is an embedding function mapping a length-*K k*-mer to a *l*-dimensional continuous representation; the embedding dimension *l* and the *k*-mer length *K* are two model hyper-parameters. Subsequently, *A* and *D* are represented by the corresponding *k*-mer embedding sequences, 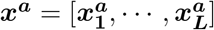 and 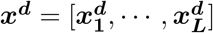.

#### Bidirectional RNNs

Based on *k*-mer embedding vectors, we apply bidirectional RNNs (BiRNNs) to learn the sequential properties in genes. The *k*-mer embedding sequences are scanned twice in both directions as forward and backward passes. During the forward pass, BiRNNs compute forward hidden states 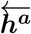 and 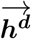 as:

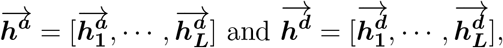

where 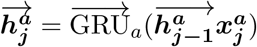; 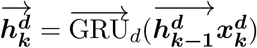. 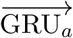 and 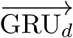 are gated recurrent units (GRUs) [8] with different parameters for acceptors and donors, respectively. Note that we adopt GRUs instead of other RNN cells like long-short term memory (LSTM) [24] because GRUs require fewer parameters [28]. Similarly, the backward pass reads the sequences in the opposite order, thereby calculating backward hidden states 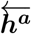 and 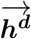 as:

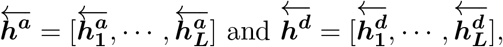

where 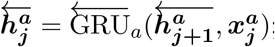; 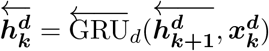. To model *k*-mers with context information, we concatenate forward and backward hidden states as the hidden representations of *k*-mers in *A* and *D* as:

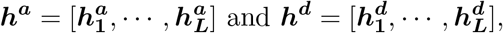

where 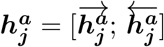; 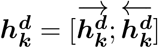.

#### *k*-mer Attention

Since different *k*-mers can have unequal importance for representing the properties of splice sites, we introduce the attention mechanism [3] to identify and aggregate the hidden representations of *k*-mers that are more important than others. More precisely, the importance scores of representations 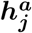 and 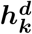 can be estimated by the *k*-mer attention vectors 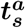 and 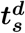 as:

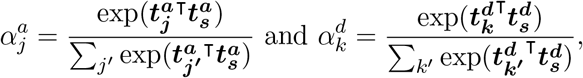

where 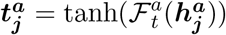; 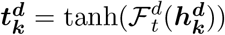; 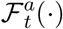 and 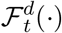 are fully-connected layers. tanh(·) is the activation function for the convenience of similarity computation. The importance scores are first measured by the inner-products to the *k*-mer attention vectors and then normalized by a softmax function over the scores of all *k*-mers. Note that the *k*-mer attention vectors 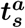 and 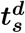 are learnable and updated during optimization as model parameters. Finally, the acceptor embedding *w*^*a*^ of *A* and the donor embedding *w*^*d*^ of *D* can be derived by aggregating the hidden representations of *k*-mers weighted by their learned importance scores as:

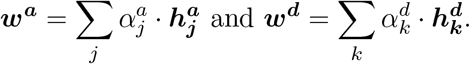

### 3.4 Cross-attention for Modeling Deep Interaction

Modeling interactions among splice sites is essential for circular RNA prediction because back-splices occur when the donors prefer the upstream acceptors over the downstream ones. Inspired by recent successes in natural language processing [22] and computer vision [32], we propose the cross-attention layer to learn deep interaction between acceptors and donors.

#### Cross-attention Layer

For acceptors, the cross-attention layer aims at deriving cross-attentive acceptor embeddings that not only represent the acceptor sites and their flanking regions but also preserve the knowledge of relevant donors from donor embeddings. Similarly, the cross-attentive donor embeddings are simultaneously obtained for donors. To directly model relations between embeddings, we adopt the dot-product attention mechanism [46] for the cross-attention layer. For each acceptor embedding 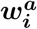 the relevance of a donor embedding 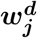 can be computed by a dot-product 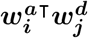 so that the attention weights 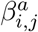 be calculated with a softmax function over all donors. Likewise, the attention weights 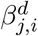 each donor embedding 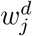 can also be measured by dot-products to the acceptor embeddings. Stated formally, we have:

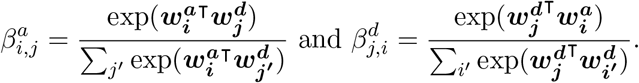

Therefore, the cross-attentive embeddings of acceptors and donors can then be derived by aggregations based on the attention weights as:

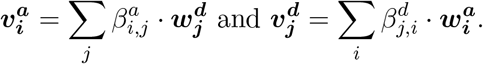

Note that we do not utilize the multi-head attention mechanism [46] because it requires much more massive training data to learn multiple projection matrices. As shown in Section 4, the vanilla dot-product attention is sufficient to obtain satisfactory predictions with significant improvements over baselines.

### 3.5 Circular RNA Prediction

To predict circRNAs, we apply the attention mechanism [3] again to aggregate cross-attentive acceptor and donor embeddings into an acceptor representation and a donor representation as ultimate features to predict circRNAs.

#### Acceptor and Donor Attention

Although the cross-attention layer provides information cross-attentive embeddings for all acceptors and donors, most of the splice sites can be irrelevant to backsplicing. To tackle this issue, we present the acceptor and donor attention to identify splice sites that are more important than other ones. Similar to *k*-mer attention, the importance scores of cross-attentive embeddings for acceptors and donors can be computed as:

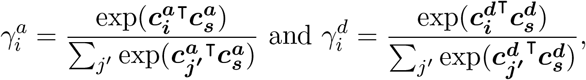

where 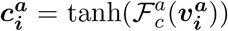; 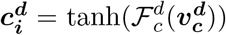; 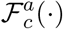 and 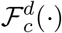 are fully-connected layers. Subsequently, the acceptor and donor representations *r*_*a*_ and *r*_*d*_ can be derived based on the attention weights of cross-attentive embeddings as:

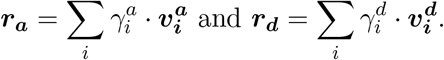

#### Prediction as Binary Classification

Here we treat circular RNA prediction as a binary classification task. More specifically, we estimate a probabilistic score *ŷ* to approximate the probability of existing a circRNA. The ultimate features *r* for machine learning are provided by concatenating the acceptor and donor representations as *r* = [*r*_*a*_; *r*_*d*_]. Finally, the probabilistic score *ŷ* can be computed by a sigmoid function with a fully-connected layer as follows:

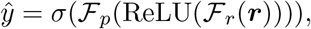

where 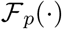 and 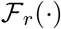 are fully-connected layers; ReLU(·) is the activation function for the hidden layer [16]; *σ*(·) is the logistic sigmoid function [18]. The binary prediction can be further generated by a binary indicator function as 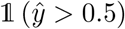.

### 3.6 Learning and Optimization

To solve circular RNA prediction as a binary classification problem, JEDI is optimized with a binary cross-entropy [23]. Formally, the loss function for optimization can be written as follows:

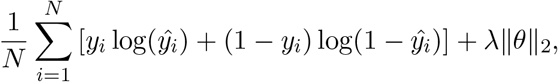

where *N* is the number of training gene sequences; *y*_*i*_ is a binary indicator demonstrating whether the *i*-th training sequence exists a circRNA; *ŷ*_*i*_ is the approximated probabilistic score for the *i*-th training gene sequence; *λ* is the L2-regularization weight for the set of model parameters *θ*.

### 3.7 Remarks on the Interpretability of JEDI

The usage of attention mechanisms is one of the most essential keys in JEDI, including the donor and acceptor attention, the cross-attention layer, and the *k*-mer attention in junction encoders. In addition to choosing important information to optimize the objective, one of the most significant benefits of using attention mechanisms is the interpretability.

#### Application: Zero-shot Backsplicing Discovery

For circRNAs, the attention weights can become interpretable hints for discovering backsplicing without training on the annotated backspliced sites. For example, when the model is optimized for accurately predicting circRNAs, the weights of donor attention are reformed to denote the important and relevant donors, which are preferred for the upstream acceptors to backsplice. In other words, the probabilistic attention weight 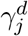 for each donor *d*_*j*_ can be interpreted as the probability of being a backsplice donor site as:

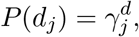

where the softmax function guarantees 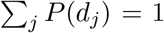. Similarly, the attention weight 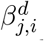 of each acceptor *a*_*i*_ for deriving the cross-attentive embedding of the donor *d*_*j*_ can be explained as the conditional probability of being selected as the backsplice acceptor site from the donor *d*_*j*_ as:

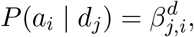

where we also have the probabilistic property 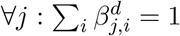 from the softmax function. Based on the above interpretations, for any pair of a donor *d*_*j*_ and an acceptor *a*_*i*_, the probability of forming a backsplice can be approximated by decomposing the joint probability *P*(*d*_*j*_, *a*_*i*_) as:

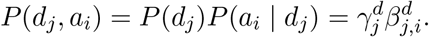

Therefore, without any training backsplice site annotation as zero-shot learning [43], we can transfer the knowledge in the training data for circular RNA prediction to discover potential backsplice sites by ranking the pairs of acceptors and donors according to *P*(*d*_*j*_, *a*_*i*_). Particularly, the interpretations can be also aligned with the process of RNA splicing, bringing more biological insights into JEDI. In Section 3.4, we further conduct experiments to demonstrate that JEDI is capable of addressing the task of zero-shot backsplicing discovery.

## 4 Experiments

In this section, we conduct extensive experiments on benchmark datasets for two tasks and in-depth analysis to verify the performance and robustness of the proposed framework, JEDI.

### 4.1 Datasets

#### Human circRNA on isoform level

We use the benchmark dataset generated by Chaabane et al. [6]. The positive data generation follows a similar setting as described in Pan and Xiong [37]. The human circRNAs are obtained from circRNADb [17], which contains 32,914 RNA molecules covering a diverse range of tissues and cell types. circRNAs with sequences shorter than 200 nucleotides are removed, resulting in 31,939 positive cases. The negative set is composed of other lncRNAs, such as processed transcripts, anti-sense, sense intronic and sense overlapping. It is constructed based on the annotation provided by GENCODE v19 [12] with strong evidence. Specifically, only the experimentally validated or manually annotated transcripts are considered, resulting in 19,683 negative cases. Each dataset is equally divided into five parts to conduct 5-fold cross validation. The sequences of all positive and negative cases are based on hg19.

#### Human circRNA on gene level

To evaluate the capability of JEDI in predicting the presence of a circRNA and the back-splicing positions in a genomic region, we construct the positive and negative sets based on gene-level annotation. For the positive set, we use BEDTools [40] to identify the overlapping genes of each circRNA from circRNADb. For each gene, we extract the exon information according to the annotation provided by GENCODE v19. In addition, we also include the exons of the overlapping circRNAs if not present in GENCODE. For the negative set, we extract the corresponding gene for each negative isoform in the benchmark dataset. We remove genes that have been selected in the positive set, and ensure that the negative set covers a wide range of exon number. The final list contains 7,777 positive and 7,000 negative genes.

#### Mouse circRNA on isoform level

The mouse circRNAs are obtained through *circbase* [15] which contains public circRNA datasets for several species reported in literature. Out of the 1,903 mouse circRNAs, we remove isoforms shorter than 200 nucleotides, resulting in 1,522 positive cases. Using the annotation provided by GENCODE vM1, we randomly select other lincRNAs longer than 200nt, generating 1,522 negative cases to create a balanced dataset. The sequences of all positive and negative cases are based on mm9.

### 4.2 Experimental Settings

#### Baseline Methods

To evaluate the performance of JEDI, we compare with eight competitive baseline methods, including circDeep [6], PredcircRNA [37], DeepCirCode [48], nRC [11], Support Vector Machines (SVM), Random Forest (RF), attentive-CNN (Att-CNN), and attentive-RNN (Att-RNN). Specifically, circDeep and PredcircRNA are the state-of-the-art circular RNA prediction methods. DeepCirCode originally takes individual splice site pairs for backsplicing prediction, which is another research problem, and leads to an enormous number of false alarms in our problem settings. To conduct fair comparisons, we modify DeepCirCode by extending the inputs to all sites and aggregating CNN representations for acceptors and donors with two max-pooling layers before applying its model structure. nRC represents lncRNA classification methods that are compatible to solve circular RNA prediction as a sequence classification problem. SVM and RF apply conventional statistical learning frameworks with the compositional *k*-mer features proposed by Wang and Wang [47] for backsplicing prediction. Attentive CNN and RNN as popular deep learning approaches utilize CNNs and RNNs with the attention mechanism [3] for sequence modeling, thereby predicting circRNAs based on a fully-connected hidden layer with the ReLU activation function [16].

#### Evaluation Metrics and Protocol

Six conventional binary classification metrics are selected as the evaluation metrics for both tasks, including the overall accuracy (Acc), precision (Prec), sensitivity (Sens), specificity (Spec), F1-score, and Matthew correlation coefficient (MCC) on positive cases. For all metrics, the higher metric scores indicate more satisfactory performance. We conduct a 5-fold cross-validation for evaluation on both isoform-level and gene-level circular RNA prediction. Specifically, for each task, the data are randomly shuffled and evenly partitioned into five non-overlapping subsets. In the five folds of experiments, each subset has a chance to be considered as the testing data for assessing the model trained by the remaining four subsets, thereby ensuring an unbiased and fair evaluation. Finally, we evaluate the methods by aggregating the scores over the 5-fold experiments for each metric.

#### Implementation Details

Our approach, JEDI, is implemented in Tensorflow [1] and released in GitHub as shown in Abstract. The AMSGrad optimizer [41] is adopted to optimize the model parameters with a learning rate *η* = 10^−3^, exponential decay rates *β*_1_ = 0.9 and *β*_2_ = 0.999, a batch size 64, and an L2-regularization weight *λ* = 10^−3^. As the hyper-parameters of JEDI, the *k*-mer size *K* and the number of dimensions *l* for *k*-mer embeddings are set to 3 and 128. We set the length of flanking regions *L* to 4. The hidden state size of GRUs for both directions in junction encoders is 128. The size of all attention vectors is set to 16. The number of units in the fully-connected hidden layer 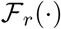 for circular RNA prediction is 128. The model parameters are trained until the convergence for each fold in cross-validation. For the baseline methods, the experiments for circDeep, PredcircRNA, and nRC are carried out according to the publicly available implementations released by the authors of original papers. SVM and RF are implemented in Python with the scikit-learn library [38]. As for deep learning approaches, DeepCirCode, Attentive-CNN, and Attentive-RNN are implemented in Tensorflow, which is the same as our proposed JEDI. For all methods, we conduct parameter fine-tuning for fair comparisons. All of the experiments are also equitably conducted on a computational server with one NVIDIA Tesla V100 GPU and one 20-core Intel Xeon CPU E5-2698 v4 @ 2.20GHz.

### 4.3 Isoform-level Circular RNA Prediction

Table 1 shows the performance of all methods for isoform-level circular RNA prediction. Among the baseline methods, circDeep as the state-of-the-art approach and DeepCirCode considering junctions perform the best. It is because circDeep explicitly accounts for the reverse complimentary sequence matches in flanking regions of the junctions, and DeepCirCode models the flanking regions with deep learning. Consistent with the previous study [6], PredcircRNA performs worse than circDeep. With compositional *k*-mer based features designed for backsplicing prediction, SVM and RF surprisingly outperform PredicrcRNA by 12.27% and 15.84% in accuracy. It not only shows that the *k*-mers are universally beneficial across different tasks but also emphasizes the rationality of using *k*-mers for junction encoders in JEDI. As an lncRNC classification method, nRC also shows its potential for circRNA prediction with a 13.18% improvement over PredcircRNA in accuracy. Although Att-CNN and Att-RNN utilize the attention mechanism, they can only model the whole sequences and present limited performance without any knowledge of junctions. As our proposed approach, JEDI significantly outperforms all of the baseline methods across all evaluation metrics. Particularly, JEDI achieves 9.89% and 7.60% improvements over DeepCirCode in accuracy and F1-score, respectively. The experimental results have demonstrated the effectiveness of junction encoders and the crossattention layer that models deep interaction among splice sites.

**Table 1:** Evaluation of isoform-level circular RNA prediction based on the 5-fold cross-validation. We report the mean and standard deviation for each metric.

| Method | Accuracy | Precision | Sensitivity | Specificity | F1-score | MCC |
| --- | --- | --- | --- | --- | --- | --- |
| SVM | $0.7386 \pm 0.0162$ | $0.7580 \pm 0.0493$ | $0.8610 \pm 0.0727$ | $0.5401 \pm 0.1542$ | $0.8027 \pm 0.0078$ | $0.4362 \pm 0.0470$ |
| RF | $0.7621 \pm 0.0025$ | $0.7773 \pm 0.0017$ | $0.8626 \pm 0.0039$ | $0.5989 \pm 0.0041$ | $0.8177 \pm 0.0022$ | $0.4833 \pm 0.0054$ |
| Att-CNN | $0.7677 \pm 0.0041$ | $0.7854 \pm 0.0064$ | $0.8595 \pm 0.0092$ | $0.6187 \pm 0.0175$ | $0.8207 \pm 0.0032$ | $0.4968 \pm 0.0094$ |
| Att-RNN | $0.7713 \pm 0.0068$ | $0.7852 \pm 0.0139$ | $0.8684 \pm 0.0231$ | $0.6136 \pm 0.0402$ | $0.8244 \pm 0.0061$ | $0.5046 \pm 0.0155$ |
| nRC | $0.7446 \pm 0.0035$ | $0.7807 \pm 0.0254$ | $0.8202 \pm 0.0499$ | $0.6220 \pm 0.0773$ | $0.7986 \pm 0.0110$ | $0.4537 \pm 0.0123$ |
| PredcircRNA | $0.6579 \pm 0.0098$ | $0.7065 \pm 0.0195$ | $0.5997 \pm 0.0095$ | $0.7224 \pm 0.0302$ | $0.6485 \pm 0.0040$ | $0.3238 \pm 0.0230$ |
| circDeep | $0.8904 \pm 0.0051$ | $0.9534 \pm 0.0133$ | $0.8326 \pm 0.0046$ | $0.9546 \pm 0.0136$ | $0.8889 \pm 0.0044$ | $0.7888 \pm 0.0119$ |
| DeepCirCode | $0.9008 \pm 0.0169$ | $0.9039 \pm 0.0350$ | $0.9418 \pm 0.0188$ | $0.8344 \pm 0.0710$ | $0.9218 \pm 0.0110$ | $0.7898 \pm 0.0354$ |
| **JEDI** | **$0.9899 \pm 0.0012$** | **$0.9926 \pm 0.0010$** | **$0.9912 \pm 0.0024$** | **$0.9880 \pm 0.0017$** | **$0.9919 \pm 0.0010$** | **$0.9787 \pm 0.0026$** |

### 4.4 Gene-level Circular RNA Prediction

We further evaluate all methods on gene-level circular RNA prediction. Note that this task is more difficult than the isoform-level prediction because each junction can be a backsplice site. Since a full gene sequence can encode for multiple isoforms, there can be multiple site pairs forming backsplices for different isoforms. Consequently, models cannot learn from absolute positions for circRNA prediction. As shown in Table 2, all methods deliver worse performance than the results in isoform-level circRNA prediction. Notably, the evaluation metrics have demonstrated a similar trend as shown in Table 1. DeepCirCode and circDeep are still the best baseline methods, showing the robustness of exploiting the knowledge about splice junctions. SVM, RF, and nRC still outperform PredicircRNA by at least 13.18% in accuracy. Att-CNN and Att-RNN using the attention mechanism still fail to obtain extraordinary performance because they are unaware of junction information, which is essential for backsplicing events. In this more difficult task, JEDI consistently surpasses all of the baseline methods across all evaluation metrics. For instance, JEDI beats DeepCirCode by 10.54% and 9.73% in accuracy and F1-score, respectively. The experimental results further reveal that our proposed JEDI is capable of tackling different scenarios of circular RNA prediction with consistently satisfactory predictions.

**Table 2:** Evaluation of gene-level circular RNA prediction based on the 5-fold cross-validation. We report the mean and standard deviation for each metric.

| Method | Accuracy | Precision | Sensitivity | Specificity | F1-score | MCC |
| --- | --- | --- | --- | --- | --- | --- |
| SVM | 0.7248 ± 0.0596 | 0.7859 ± 0.1163 | 0.7242 ± 0.1707 | 0.7256 ± 0.2859 | 0.7324 ± 0.0403 | 0.4833 ± 0.0898 |
| RF | 0.7255 ± 0.0088 | 0.6985 ± 0.0056 | 0.8416 ± 0.0159 | 0.5964 ± 0.0089 | 0.7634 ± 0.0089 | 0.4542 ± 0.0193 |
| Att-CNN | 0.7387 ± 0.0078 | 0.7098 ± 0.0157 | 0.8534 ± 0.0263 | 0.6113 ± 0.0403 | 0.7746 ± 0.0051 | 0.4824 ± 0.0136 |
| Att-RNN | 0.7486 ± 0.0158 | 0.7275 ± 0.0256 | 0.8580 ± 0.0415 | 0.6112 ± 0.0721 | 0.7869 ± 0.0246 | 0.4914 ± 0.0217 |
| nRC | 0.7293 ± 0.0131 | 0.7440 ± 0.0216 | 0.7428 ± 0.0465 | 0.7143 ± 0.0474 | 0.7424 ± 0.0178 | 0.4587 ± 0.0262 |
| PredcircRNA | 0.6250 ± 0.0125 | 0.6656 ± 0.0206 | 0.5795 ± 0.0090 | 0.6754 ± 0.0355 | 0.6193 ± 0.0047 | 0.2558 ± 0.0280 |
| circDeep | 0.8493 ± 0.0042 | 0.8888 ± 0.0146 | 0.8161 ± 0.0090 | 0.8861 ± 0.0177 | 0.8507 ± 0.0027 | 0.7018 ± 0.0104 |
| DeepCirCode | 0.8726 ± 0.0134 | 0.8696 ± 0.0242 | 0.8934 ± 0.0427 | 0.8496 ± 0.0372 | 0.8805 ± 0.0152 | 0.7463 ± 0.0257 |
| **JEDI** | **0.9646 ± 0.0014** | **0.9703 ± 0.0049** | **0.9622 ± 0.0056** | **0.9673 ± 0.0057** | **0.9662 ± 0.0014** | **0.9291 ± 0.0027** |

### 4.5 Independent Study on Mouse circRNAs

To demonstrate the robustness of JEDI, we conduct an independent study on the dataset of mouse circRNAs. Previous studies have shown that circRNAs are evolutionarily conserved [4, 27], and thus we evaluate the potential of predicting the circRNAs across different species. More precisely, we train each method using the human dataset on isoform-level, thereby predicting the circuRNAs on the mouse dataset. Note that some of the required features for PredcircRNA are missing on the mouse datasets. In addition to this, PredictcRNA perform the worst in other experiments. For these reasons, we exclude PredcircRNA from this study. Table 3 presents the experimental results of the independent study. Compared to the experiments conducted on the same species as shown in Table 1, most of the deep learning methods have slightly lower performance because they are specifically optimized for human data; SVM and RF have similar performance in the independent study probably because *k*-mer features are simpler and more general to different species. Interestingly, the accuracy of circDeep significantly drops in the study. It is likely due to the fact that circDeep heavily pre-trains the sequence modeling on human data with the serious over-fitting phenomenon. As a result, our proposed JEDI still outperforms all of the baseline methods. It demonstrates that JEDI is robust across the datasets of different species.

**Table 3:**
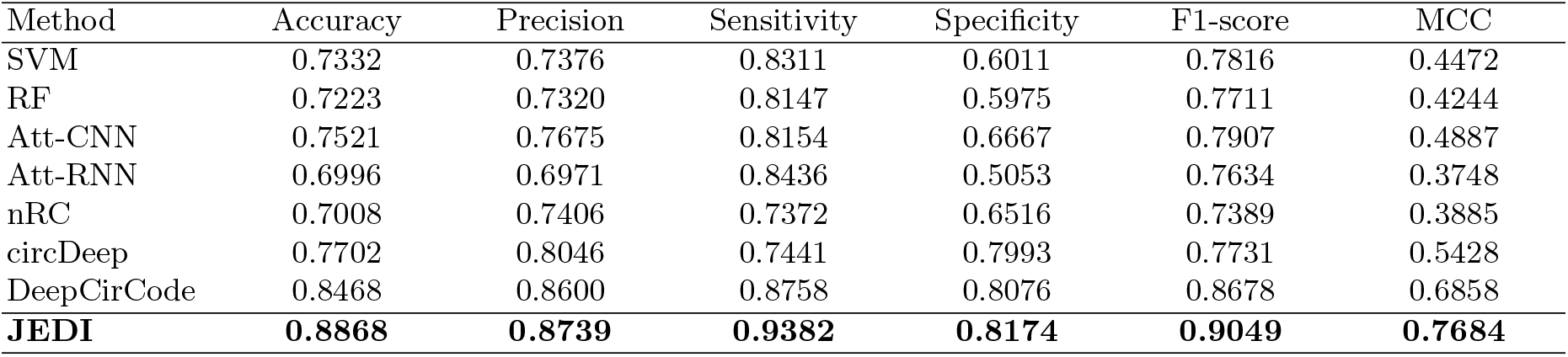
Independent study of isoform-level circular RNA prediction for mouse circRNAs based on the models trained on human circRNAs.

| Method | Accuracy | Precision | Sensitivity | Specificity | F1-score | MCC |
| --- | --- | --- | --- | --- | --- | --- |
| SVM | 0.7332 | 0.7376 | 0.8311 | 0.6011 | 0.7816 | 0.4472 |
| RF | 0.7223 | 0.7320 | 0.8147 | 0.5975 | 0.7711 | 0.4244 |
| Att-CNN | 0.7521 | 0.7675 | 0.8154 | 0.6667 | 0.7907 | 0.4887 |
| Att-RNN | 0.6996 | 0.6971 | 0.8436 | 0.5053 | 0.7634 | 0.3748 |
| nRC | 0.7008 | 0.7406 | 0.7372 | 0.6516 | 0.7389 | 0.3885 |
| circDeep | 0.7702 | 0.8046 | 0.7441 | 0.7993 | 0.7731 | 0.5428 |
| DeepCirCode | 0.8468 | 0.8600 | 0.8758 | 0.8076 | 0.8678 | 0.6858 |
| **JEDI** | **0.8868** | **0.8739** | **0.9382** | **0.8174** | **0.9049** | **0.7684** |

### 4.6 Zero-shot Backsplicing Discovery

As mentioned in Section 3.7, the interpretability of the attention mechanisms and the cross-attention layer enables JEDI to achieve zero-shot backsplicing discovery. To evaluate the performance of zeroshot backsplicing, we compute the probabilistic score *P*(*d*_*j*_, *a*_*i*_) using the attention weights 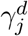 and 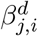, thereby indicating the likelihood of forming a backsplice for each pair of a candidate donor *d*_*j*_ and a candidate acceptor *a*_*i*_. Hence, we can simply evaluate the probabilistic scores with the receiver operating characteristic (ROC) curve and the area under the ROC curve (AUC). Note that here we still apply 5-fold cross-validation for experiments based on the gene-level human circRNA dataset. Since none of the existing methods can address the task of zero-shot backsplicing prediction, we compare with random guessing, which is equivalent to the chance line in ROCs with an AUC score of 0.5. Figure 3 depicts the ROC curves with AUC scores over five folds of experiments. The results show that the backspliced site pairs discovered by JEDI are effective with an average AUC score of 0.7992. In addition, JEDI is also robust in this task with a small standard deviation of AUC scores.

**Fig. 3:**
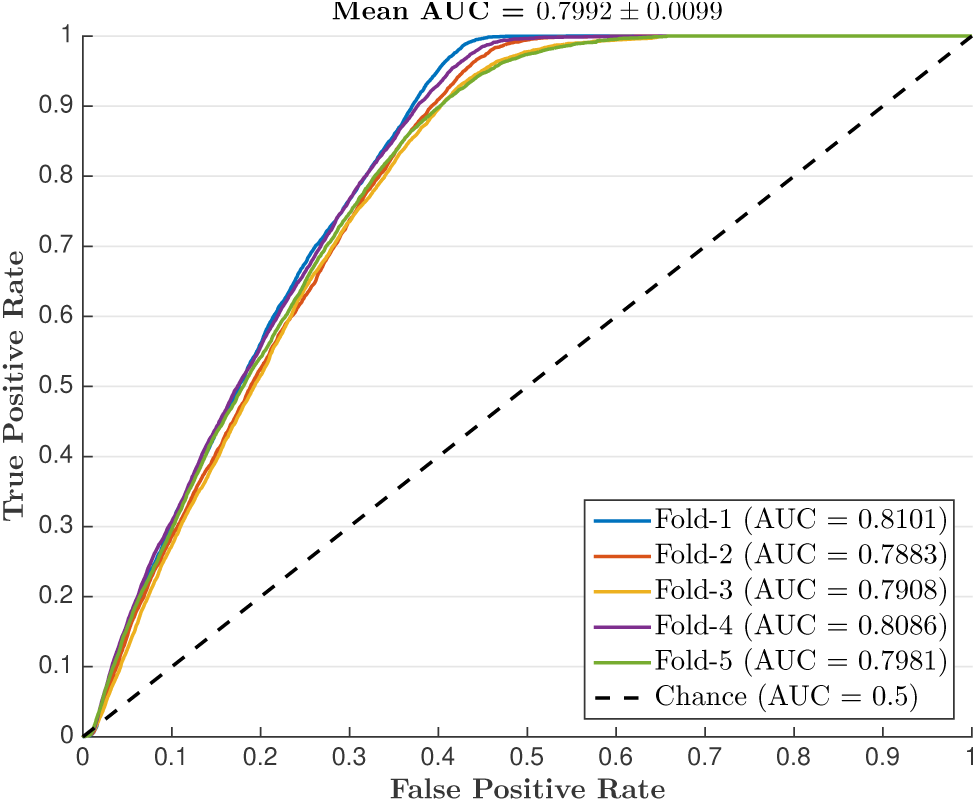
The ROC curves for zero-shot backsplicing discovery based on the 5-fold cross-validation and JEDI trained for gene-level circular RNA prediction.

### 4.7 Analysis and Discussions

In this section, we first discuss the impacts of hyper-parameters for JEDI and then conduct the run-time analysis for all methods to demonstrate verify the model efficiency of JEDI.

#### Length of Flanking Regions *L*

The flanking region length *L* for junction encoders plays an important role in JEDI to represent splice sites. Figure 4a illustrates the circular RNA prediction performance of JEDI over different flanking region lengths. For all evaluation metrics, the performance slightly improves when *L* increases to 4. However, the performance significantly drops when *L* ≥ 32. It shows that nucleotides nearer to junctions are more important than other ones for predicting backsplicing. This result is also consistent with previous studies on RNA splicing [36]. Moreover, circRNAs tend to contain fewer nucleotides than other transcripts from the same gene [26], so excessive and redundant information could only lead to noises and lower the prediction performance.

**Fig. 4:**
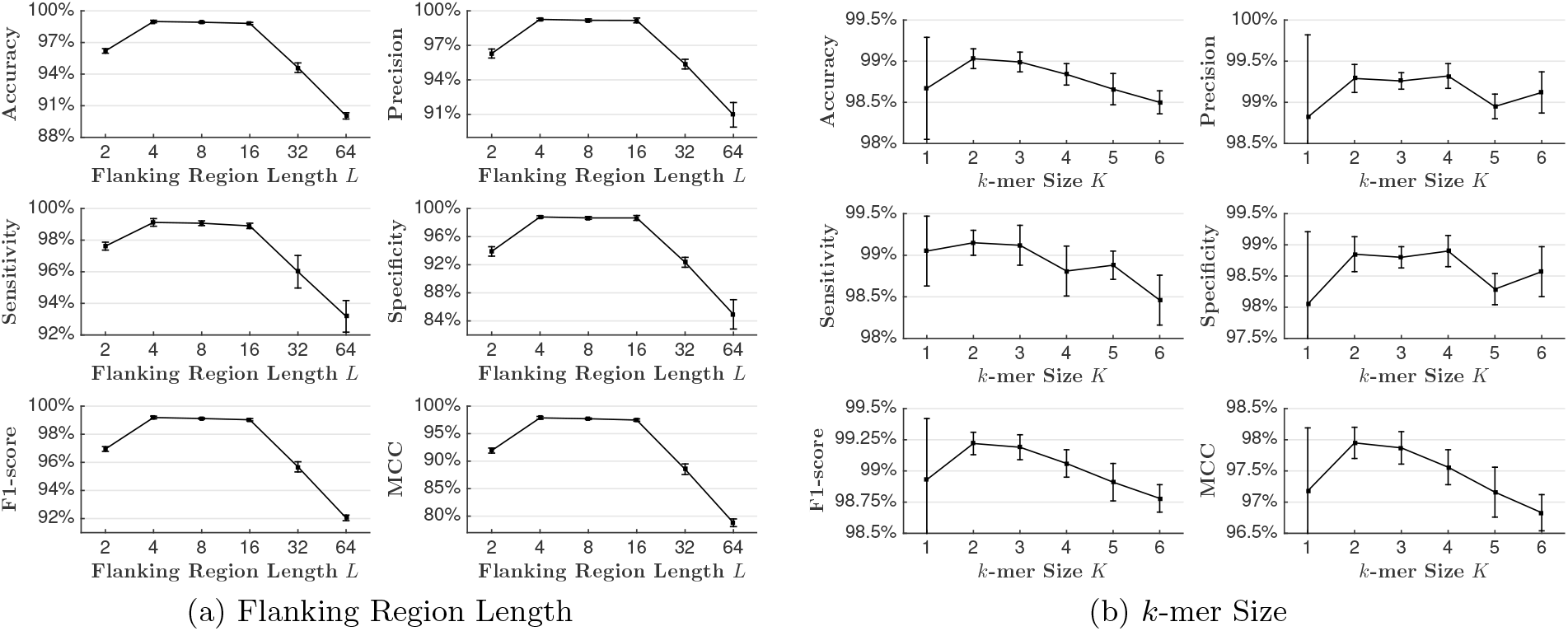
The isoform-level circular RNA prediction performance of JEDI with different flanking region lengths *L* and *k*-mer sizes *K* based on the 5-fold cross validation. We report the mean for each metric and apply error bars to indicate standard deviations.

#### Size of *k*-mers *K*

The derivation of *k*-mers is crucial for JEDI because JEDI treats *k*-mers as the fundamental inputs over gene sequences. Figure 4b shows how the size of *k*-mers affects the prediction performance. JEDI performs the best with 2-mers and 3-mers when the performance gets worse with longer or shorter *k*-mers. It could be because a small *k*-mer size makes *k*-mers less significant for representations. In addition, the embedding space of long *k*-mers could be too enormous for JEDI to learn with limited training data. It is also worthwhile to mention that 1-mers lead to much higher standard deviations because of their low significance induces high instability and sensitive embeddings during the learning process. This finding is also consistent with previous studies [42].

#### Run-time Analysis

To verify the efficiency of JEDI, we conduct the run-time analysis for all methods in our experiments based on the task of isoform-level circular RNA prediction. Note that we only consider the time in training and testing. The run-time of feature extraction and disk I/O are ignored because the features can be pre-processed. Disk I/O can be affected by many factors that are irrelevant to methods, such as I/O scheduling in operating systems. As shown in Table 4, JEDI is efficient and averagely needs only less than three minutes because it only focuses on junctions and flanking regions. Similarly, DeepCirCode, which is also a junction based deep learning method, has comparable execution time to JEDI. In contrast, Att-CNN and Att-RNN are relatively inefficient because they scan the whole sequences in every training batch, where Att-RNN with non-parallelizable recurrent units is slower. Although nRC reads the whole sequences, it runs faster than some attention-based methods because of its simpler model structure. SVM, RF, and PredcircRNA are the most efficient because they apply straightforward statistical machine learning frameworks for training. As a side note, the feature extraction of PredcircRNA is extremely expensive in execution time and averagely costs more than 28 hours to extract multi-facet features in our experiments. circDeep is the most inefficient in our experiments because it consists of many time-consuming components, such as embedding and LSTM pre-training.

**Table 4:** Run-time analysis on isoform-level circular RNA prediction in seconds (`s`), minutes (`m`), and hours (`h`), based on the 5-fold cross-validation. We report the mean of the training time (over five folds).

| Method | Time | Method | Time | Method | Time |
| --- | --- | --- | --- | --- | --- |
| SVM | 28.76s | Att-CNN | 13.35m | circDeep | >24h |
| RF | 21.03s | Att-RNN | 51.53m | DeepCirCode | 3.80m |
| nRC | 4.07m | PredcircRNA | 43.66s | JEDI | 2.75m |

## 5 Conclusions

In this paper, we propose a novel end-to-end deep learning approach for circular RNA prediction by learning to appropriately model splice sites with flanking regions around junctions and studying the deep relationships among these sites. The effective attentive junction encoders are first presented to represent each splice site when the innovative cross-attention layer is proposed to learn deep interaction among the sites. Moreover, JEDI is capable of discovering backspliced site pairs without training on annotated site pairs. The experimental results demonstrate that JEDI is effective and robust in circular RNA prediction across different data levels and the datasets of different species. The backspliced site pairs discovered by JEDI are also promising to form circular RNAs through backsplicing. The reasons and insights can be concluded as follows: (1) JEDI only models valuable and essential flanking regions around the junctions of splice sites, thereby discarding irrelevant and redundant information for circular RNA prediction. (2) the properties of splice sites and essential information for forming circular RNAs can be well-preserved by junction encoders; (3) the attention mechanisms and the cross-attention layer provide intuitive and interpretable hints to implicitly model backsplicing as demonstrated in the experiments.

## Notes

#### Summary of Updates

Update author information and fix a typo.

## Bibliography

[1] Martín Abadi, Paul Barham, Jianmin Chen, Zhifeng Chen, Andy Davis, Jeffrey Dean, Matthieu Devin, Sanjay Ghemawat, Geoffrey Irving, Michael Isard, et al. Tensorflow: A system for large-scale machine learning. In 12th {USENIX} Symposium on Operating Systems Design and Implementation ({OSDI} 16), pages 265–283, (2016).

[2] Reut Ashwal-Fluss, Markus Meyer, Nagarjuna Reddy Pamudurti, Andranik Ivanov, Osnat Bartok, Mor Hanan, Naveh Evantal, Sebastian Memczak, Nikolaus Rajewsky, and Sebastian Kadener. circrna biogenesis competes with pre-mrna splicing. Molecular cell, 56(1):55–66, (2014).

[3] Dzmitry Bahdanau, Kyunghyun Cho, and Yoshua Bengio Neural machine translation by jointly learning to align and translate. In 3rd International Conference on Learning Representations, ICLR 2015, (2015).

[4] Steven P Barrett and Julia Salzman. Circular rnas: analysis, expression and potential functions. Development, 143(11):1838–1847, (2016).

[5] Marcel Boss and Christoph Arenz. A fast and easy method for specific detection of circular rna by rolling-circle amplification. ChemBioChem, (2019).

[6] Mohamed Chaabane, Robert M Williams, Austin T Stephens, and Juw Won Park. circdeep: deep learning approach for circular rna classification from other long non-coding rna. Bioinformatics, 36(1):73–80, (2020).

[7] Lei Chen, Yu-Hang Zhang, Guohua Huang, Xiaoyong Pan, ShaoPeng Wang, Tao Huang, and Yu-Dong Cai. Discriminating cirrnas from other lncrnas using a hierarchical extreme learning machine (h-elm) algorithm with feature selection. Molecular genetics and genomics, 293(1): 137–149, (2018).

[8] Kyunghyun Cho, Bart Van Merriënboer, Caglar Gulcehre, Dzmitry Bahdanau, Fethi Bougares, Holger Schwenk, and Yoshua Bengio. Learning phrase representations using rnn encoder-decoder for statistical machine translation. arXiv preprint arXiv:1406.1078, (2014).

[9] Umber Dube, Jorge L Del-Aguila, Zeran Li, John P Budde, Shan Jiang, Simon Hsu, Laura Ibanez, Maria Victoria Fernandez, Fabiana Farias, Joanne Norton, et al. An atlas of cortical circular rna expression in alzheimer disease brains demonstrates clinical and pathological associations. Nature neuroscience, 22(11):1903–1912, (2019).

[10] Robert A Dubin, Manija A Kazmi, and Harry Ostrer. Inverted repeats are necessary for circularization of the mouse testis sry transcript. Gene, 167(1-2):245–248, (1995).

[11] Antonino Fiannaca, Massimo La Rosa, Laura La Paglia, Riccardo Rizzo, and Alfonso Urso. nrc: non-coding rna classifier based on structural features. BioData mining, 10(1):27, (2017).

[12] Adam Frankish, Mark Diekhans, Anne-Maud Ferreira, Rory Johnson, Irwin Jungreis, Jane Loveland, Jonathan M Mudge, Cristina Sisu, James Wright, Joel Armstrong, et al. Gencode reference annotation for the human and mouse genomes. Nucleic acids research, 47(D1):D766–D773, (2019).

[13] Yuan Gao, Jinfeng Wang, and Fangqing Zhao. Ciri: an efficient and unbiased algorithm for de novo circular rna identification. Genome biology, 16(1):4, (2015).

[14] Yuan Gao, Jinyang Zhang, and Fangqing Zhao. Circular rna identification based on multiple seed matching. Briefings in bioinformatics, 19(5):803–810, (2018).

[15] Petar Glažar, Panagiotis Papavasileiou, and Nikolaus Rajewsky. circbase: a database for circular rnas. Rna, 20(11):1666–1670, (2014).

[16] Xavier Glorot, Antoine Bordes, and Yoshua Bengio. Deep sparse rectifier neural networks. In Proceedings of the fourteenth international conference on artificial intelligence and statistics, pages 315–323, (2011).

[17] Mark A Hall Correlation-based feature selection of discrete and numeric class machine learning. In Proceedings of the Seventeenth International Conference on Machine Learning, pages 359—-366, (2000).

[18] Jun Han and Claudio Moraga The influence of the sigmoid function parameters on the speed of backpropagation learning. In International Workshop on Artificial Neural Networks, pages 195–201. Springer, (1995).

[19] Siyu Han, Yanchun Liang, Qin Ma, Yangyi Xu, Yu Zhang, Wei Du, Cankun Wang, and Ying Li. Lncfinder: an integrated platform for long non-coding rna identification utilizing sequence intrinsic composition, structural information and physicochemical property. Briefings in bioinformatics, 20(6):2009–2027, (2019).

[20] Thomas B Hansen, Trine I Jensen, Bettina H Clausen, Jesper B Bramsen, Bente Finsen, Christian K Damgaard, and Jørgen Kjems. Natural rna circles function as efficient microrna sponges. Nature, 495(7441):384–388, (2013).

[21] Thomas B Hansen, Morten T Venø, Christian K Damgaard, and Jørgen Kjems. Comparison of circular rna prediction tools. Nucleic acids research, 44(6):e58–e58, (2016).

[22] Yanchao Hao, Yuanzhe Zhang, Kang Liu, Shizhu He, Zhanyi Liu, Hua Wu, and Jun Zhao An end-to-end model for question answering over knowledge base with cross-attention combining global knowledge. In Proceedings of the 55th Annual Meeting of the Association for Computational Linguistics (Volume 1: Long Papers), pages 221–231, (2017).

[23] Geoffrey E Hinton and Ruslan R Salakhutdinov. Reducing the dimensionality of data with neural networks. science, 313(5786):504–507, (2006).

[24] Sepp Hochreiter and Jürgen Schmidhuber. Long short-term memory. Neural computation, 9 (8):1735–1780, (1997).

[25] Andranik Ivanov, Sebastian Memczak, Emanuel Wyler, Francesca Torti, Hagit T Porath, Marta R Orejuela, Michael Piechotta, Erez Y Levanon, Markus Landthaler, Christoph Dieterich, et al. Analysis of intron sequences reveals hallmarks of circular rna biogenesis in animals. Cell reports, 10(2):170–177, (2015).

[26] William R Jeck and Norman E Sharpless. Detecting and characterizing circular rnas. Nature biotechnology, 32(5):453, (2014).

[27] William R Jeck, Jessica A Sorrentino, Kai Wang, Michael K Slevin, Christin E Burd, Jinze Liu, William F Marzluff, and Norman E Sharpless. Circular rnas are abundant, conserved, and associated with alu repeats. Rna, 19(2):141–157, (2013).

[28] Rafal Jozefowicz, Wojciech Zaremba, and Ilya Sutskever An empirical exploration of recurrent network architectures. In ICML’15, pages 2342–2350, (2015).

[29] Chelsea Jui-Ting Ju, Jyun-Yu Jiang, Ruirui Li, Zeyu Li, and Wei Wang. Tahcoroll: An efficient approach for signature profiling in genomic data through variable-length k-mers. bioRxiv, page 229708, (2017).

[30] Erika Lasda and Roy Parker. Circular rnas: diversity of form and function. Rna, 20(12): 1829–1842, (2014).

[31] Yann LeCun, Yoshua Bengio, and Geoffrey Hinton. Deep learning. nature, 521(7553):436–444, (2015).

[32] Kuang-Huei Lee, Xi Chen, Gang Hua, Houdong Hu, and Xiaodong He Stacked cross attention for image-text matching. In Proceedings of the European Conference on Computer Vision (ECCV), pages 201–216, (2018).

[33] Zhaoyong Li, Chuan Huang, Chun Bao, Liang Chen, Mei Lin, Xiaolin Wang, Guolin Zhong, Bin Yu, Wanchen Hu, Limin Dai, et al. Exon-intron circular rnas regulate transcription in the nucleus. Nature structural & molecular biology, 22(3):256, (2015).

[34] Sebastian Memczak, Marvin Jens, Antigoni Elefsinioti, Francesca Torti, Janna Krueger, Agnieszka Rybak, Luisa Maier, Sebastian D Mackowiak, Lea H Gregersen, Mathias Munschauer, et al. Circular rnas are a large class of animal rnas with regulatory potency. Nature, 495(7441): 333–338, (2013).

[35] Xu Min, Wanwen Zeng, Ning Chen, Ting Chen, and Rui Jiang. Chromatin accessibility prediction via convolutional long short-term memory networks with k-mer embedding. Bioinformatics, 33(14):i92–i101, (2017).

[36] Yasumi Ohshima and Yoshie Gotoh. Signals for the selection of a splice site in pre-mrna: computer analysis of splice junction sequences and like sequences. Journal of molecular biology, 195(2):247–259, (1987).

[37] Xiaoyong Pan and Kai Xiong. Predcircrna: computational classification of circular rna from other long non-coding rna using hybrid features. Molecular Biosystems, 11(8):2219–2226, (2015).

[38] Fabian Pedregosa, Gaël Varoquaux, Alexandre Gramfort, Vincent Michel, Bertrand Thirion, Olivier Grisel, Mathieu Blondel, Peter Prettenhofer, Ron Weiss, Vincent Dubourg, et al. Scikitlearn: Machine learning in python. Journal of machine learning research, 12(Oct):2825–2830, (2011).

[39] Shibin Qu, Zhengcai Liu, Xisheng Yang, Jingshi Zhou, Hengchao Yu, Rui Zhang, and Haimin Li. The emerging functions and roles of circular rnas in cancer. Cancer letters, 414:301–309, (2018).

[40] Aaron R Quinlan and Ira M Hall. Bedtools: a flexible suite of utilities for comparing genomic features. Bioinformatics, 26(6):841–842, (2010).

[41] Sashank J Reddi, Satyen Kale, and Sanjiv Kumar On the convergence of adam and beyond. In The Sixth International Conference on Learning Representations (ICLR), (2018).

[42] Ahmet Sacan and I Hakki Toroslu Approximate similarity search in genomic sequence databases using landmark-guided embedding. In First International Workshop on Similarity Search and Applications (sisap 2008), pages 43–50. IEEE, (2008).

[43] Richard Socher, Milind Ganjoo, Christopher D Manning, and Andrew Ng Zero-shot learning through cross-modal transfer. In Advances in neural information processing systems, pages 935–943, (2013).

[44] Linda Szabo, Robert Morey, Nathan J Palpant, Peter L Wang, Nastaran Afari, Chuan Jiang, Mana M Parast, Charles E Murry, Louise C Laurent, and Julia Salzman. Statistically based splicing detection reveals neural enrichment and tissue-specific induction of circular rna during human fetal development. Genome biology, 16(1):126, (2015).

[45] Laurent F Thomas and Pal Satrom. Circular rnas are depleted of polymorphisms at microrna binding sites. Bioinformatics, 30(16):2243–2246, (2014).

[46] Ashish Vaswani, Noam Shazeer, Niki Parmar, Jakob Uszkoreit, Llion Jones, Aidan N Gomez, Lukasz Kaiser, and Illia Polosukhin Attention is all you need. In Advances in neural information processing systems, pages 5998–6008, (2017).

[47] Jun Wang and Liangjiang Wang Prediction of back-splicing sites reveals sequence compositional features of human circular rnas. In 2017 IEEE 7th International Conference on Computational Advances in Bio and Medical Sciences (ICCABS), pages 1–6. IEEE, (2017).

[48] Jun Wang and Liangjiang Wang. Deep learning of the back-splicing code for circular rna formation. Bioinformatics, 35(24):5235–5242, (2019).

[49] Kai Wang, Darshan Singh, Zheng Zeng, Stephen J Coleman, Yan Huang, Gleb L Savich, Xiaping He, Piotr Mieczkowski, Sara A Grimm, Charles M Perou, et al. Mapsplice: accurate mapping of rna-seq reads for splice junction discovery. Nucleic acids research, 38(18):e178–e178, (2010).

[50] Chun-Ying Yu and Hung-Chih Kuo. The emerging roles and functions of circular rnas and their generation. Journal of biomedical science, 26(1):29, (2019).

[51] Xiao-Ou Zhang, Hai-Bin Wang, Yang Zhang, Xuhua Lu, Ling-Ling Chen, and Li Yang. Complementary sequence-mediated exon circularization. Cell, 159(1):134–147, (2014).

